# Sex differences in the strength of selection under facultative sex

**DOI:** 10.1101/2021.10.21.465271

**Authors:** Thomas R. Haaland, Dieter Ebert, Hanna Kokko

## Abstract

Facultative sex, the ability to reproduce both sexually and asexually, is widespread across the tree of life. In anisogamous species, the frequency of sex modulates selection on traits with sex-specific expression. Current theory on conditional gene expression posits that the strength of selection on loci only expressed by a subset of individuals, and/or in a subset of environments, is proportional to the frequency of expressers in the population. We show here that this assumption does not hold when the subsets in question are males or females (because of the Fisher condition) and is most important in facultatively sexual populations. In this case, the proportion of sexually produced offspring is not determined by male frequency (sex ratio), but on relative female investment in sexual versus asexual reproduction. This breaks the link between the frequency of expressers and selection for loci with sex-specific expression. However, certain conditions can re-establish this link, for example male traits that predict mating success better under strong male-male competition, and sex ratio affecting the relative fecundity of sexual versus asexual females. Our work highlights the importance of the Fisher condition for understanding the efficiency of selection and has implications for differences in the genetic load among sexes.

## Introduction

Natural selection is not always an efficient evolutionary process. When fitness effects of mutations are small, beneficial mutations may be lost from populations by chance, and genes may succumb to degradation from slightly deleterious mutations (Kimura 1986). This can have marked consequences for selection on conditionally expressed loci, i.e. loci that are only expressed by a subset of individuals and/or in a subset of environments, because the overall selection coefficient is reduced by individuals in which the polymorphism has no fitness consequences (Van Tienderen 1991; Linksvayer and Wade 2009). All else being equal, selection can be assumed to become weakened in proportion to the frequency of gene expression (Van Dyken and Wade 2010). Similar reasoning holds for traits that are selectively neutral (or nearly neutral) under most conditions, but beneficial under rare environmental conditions (Kawecki 1994). Thus, both the origin and maintenance of traits that help an organism cope with specific (and potentially rare) conditions is a general problem in evolution.

Arguably the most common situation of conditional gene expression is related to sex-linked expression, with thousands of genes being reported to have sex-linked expression (Parsch and Ellegren 2013; Mank 2017; Molinier et al. 2018). Conditional expression can be even more extreme when sex-linked gene expression occurs in facultatively sexual species, i.e. species that are capable of both sexual and asexual reproduction. In facultative sexuals, the frequency of sexual reproduction can range from almost obligate (with parthenogenesis only as a worst-case option, as in many facultatively sexual reptiles; Watts et al. 2006; Kearney et al. 2009) to only occurring once every several thousand generations (e.g. some sex occurs even in lineages long thought to be entirely asexual, such as bdelloid rotifers; Vakhrusheva et al. 2020).

Many facultatively sexual animals exhibit cyclical parthenogenesis, a reproductive mode in which sexual reproduction occurs after one or multiple generations of clonal reproduction. In cyclical parthenogens such as aphids (Simon et al. 2002), rotifers (Smith and Snell 2012) and *Daphnia* (Decaestecker et al. 2009), sexual reproduction is typically associated with indicators for unfavorable conditions, such as the end of the growth season, winter period or dry season. It involves asexually reproducing females to produce male offspring and sexual daughters, and often leads to production of morphs or structures that are better able to cope with the changing conditions, such as diapause structures (Ram and Hadany 2016; Gerber and Kokko 2018). Sex-specific traits in facultatively sexual systems therefore provide an intriguing arena in which to study selection on and maintenance of rarely expressed traits. Any traits with male-specific functions belong to this category, especially since males often remain rare in cyclical parthenogens even in sexual generations. Both empirical and theoretical research suggests that rare sexual reproduction, and particular in combination with rarity of males, may substantially weaken the average strength of selection acting on male-specific traits (Chasnov and Chow 2002; Brisson and Nuzhdin 2008; Van Dyken and Wade 2010). However, as we show here, this effect is much less strong than previously believed.

Van Dyken and Wade (2010) developed a general theory for selection on a trait that is only expressed in some generations and/or by only a subset of individuals within a generation. Their model is, in principle, applicable to a wide range of situations, e.g., constraints on the evolution of plasticity to rarely experienced environments (Whitlock 1996; Kawecki et al. 1997; Snell-Rood et al. 2010), or selection on kin-selected genes (Linksvayer and Wade 2009; Van Dyken et al. 2011). The key result of their model is that the efficiency of selection is weakened substantially if a gene is only rarely expressed, and that such genes therefore accumulate deleterious mutations much faster than unconditionally expressed genes. Van Dyken and Wade (2010) also discuss the results of their model in the context of facultative sex, including cases where genes are expressed only in males. However, they here make the following tacit assumption: If males form a proportion *Φ* of a given generation, then selection on a male-specific locus can be derived identically to a model where the proportion *Φ* refers to any subset of all individuals, where this subset is *a priori* responsible for a proportion *Φ* of offspring produced in that generation (by ‘a priori’ we refer to the fact that differential success of individuals with different genotypes at the locus in question is not yet taken into account).

Here we argue that this assumption of Van Dyken and Wade (2010) ceases to be a valid when the genes in question are only expressed in males, because males cannot reproduce independently of females. Consider a facultatively sexual population where only a subset of females are sexual in a given generation (the rest reproduce asexually). Every offspring produced has a mother, but only sexual offspring have a father. Assuming no fecundity differences between modes of reproduction (an assumption we also relax later), the probability that a newly formed offspring has a father only depends on the proportion of offspring that were produced sexually – not on the number of males. Thus, the subset of reproductive events where male traits matter, and are exposed to selection, is in this simple scenario independent of population sex ratio.

Our aim is to complement existing population genetic theory (Fig. 1, nodes 1-2) by extending selection models with conditional gene expression (Van Dyken and Wade 2010) to sex-limited expression when sex is not obligate (Fig. 1, nodes 3-5). We investigate the reduction in the efficiency of selection on sex-specific traits and its evolutionary consequences by modeling the change in allele frequencies at a single locus, and compare it to loci expressed in every individual (constitutively expressed loci). In addition to sex specificity (Fig. 1, node 4), we compare results where the locus is expressed in females irrespective of reproductive mode with an alternative setting where the relevant trait is only expressed in sexual females (Fig. 1, node 5). Traits that matter for sexual females in contexts of mate finding, mate choice or the production of fertilizable eggs (should they differ from asexual eggs) are all examples of the latter kind (Otto 2009; Lehtonen et al. 2012). Note however that genes on the Y (or W) chromosome are not affected by these considerations, being only expressed in the heterogametic sex. In addition to our baseline model presented in the main text, we also show (Supplementary Material, section 2) that, under certain additional assumptions, a link between sex ratio (frequency of expressers) and the strength of selection on sex-limited traits can be re-established, although not one of strict proportionality.

**Figure 1:**
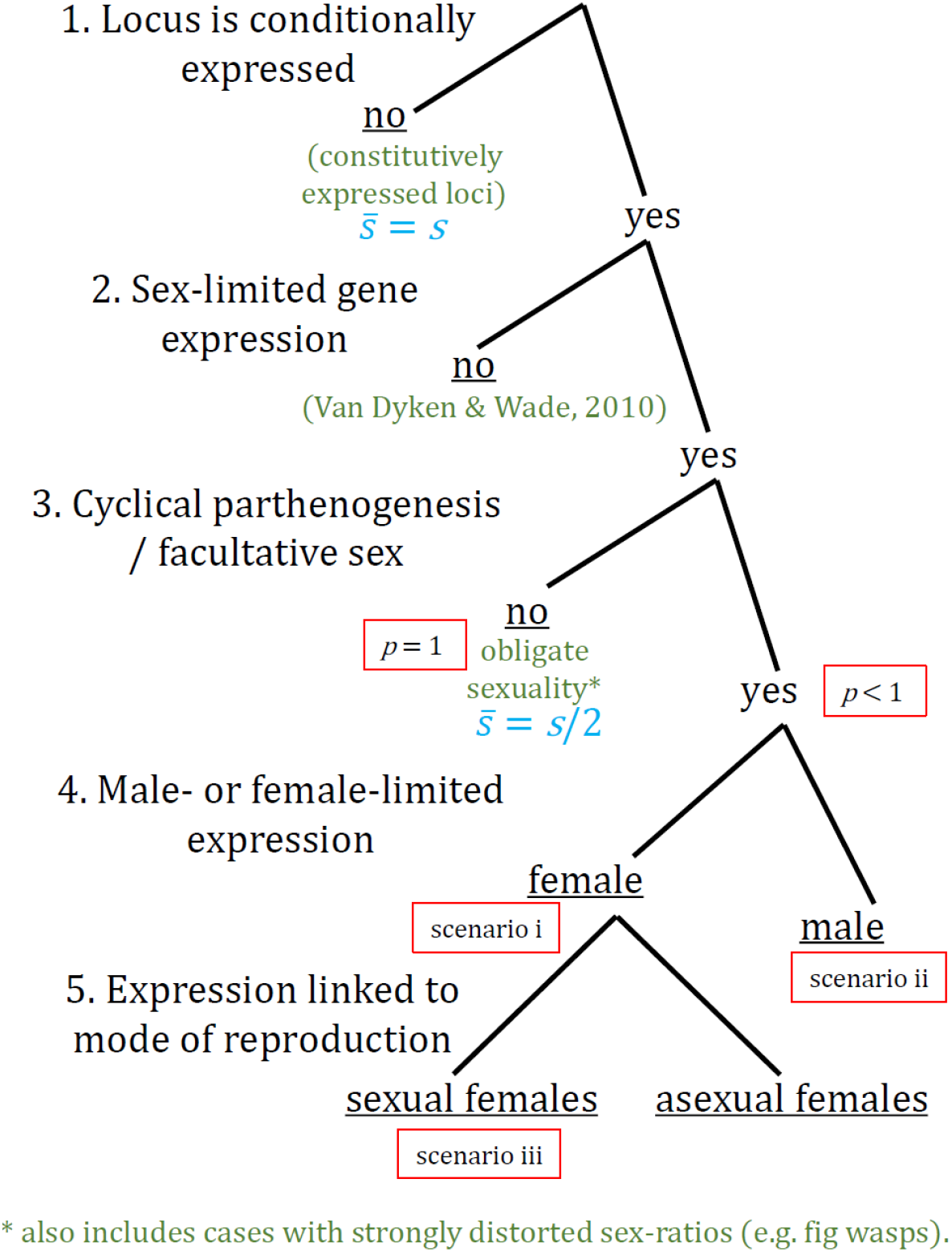
Overview of how the current paper fits into a wider classification of selection models with conditional gene expression. Colored text: Appropriate existing modeling approaches (green) with results for how selection coefficients *s* relate to population-wide realized selection 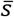 (blue). Underlined responses: Typical end points of interest with the foci of our model analyses inside red boxes (*p* indicates the frequency of sexual reproduction).

## Model description

We assume a facultatively sexual population and focus on one haploid, biallelic locus L with alleles *A* (wild type) and *a* (mutant). The *a* allele’s frequency in the parental population is denoted *x*, with no need to distinguish between females and males since we assume that the L locus impacts neither the probability of sexual reproduction nor the sex ratio. However, selection acting on locus L can differ depending on the sex and reproductive mode of the expresser. We denote the selection coefficient acting on allele *a* relative to allele *A* as *s*_m_ if expressed in males, *s*_f_ if expressed in sexual females, and *s*_a_ if expressed in asexual females. These selection coefficients can be positive (allele *a* beneficial), zero (no selection on the locus) or negative (allele *a* deleterious), with –1 denoting lethal effects. We here consider three main scenarios. i) Locus L expressed in all females (*s*_a_ = *s*_f_ ≠ 0, *s*_m_ = 0); ii) Locus L only expressed in males (*s*_a_ = *s*_f_ = 0, *s*_m_ ≠ 0); iii) Locus L only expressed in sexual females (*s*_f_ ≠ 0, *s*_a_ = *s*_m_ = 0).

### Selection over one sexual generation

In a partly sexual generation (at time *t*), the parental population consists of a proportion *q* of males and 1–*q* females. A proportion *p* of offspring is produced sexually, the rest (1–*p*) asexually. *p* is therefore interpretable as either the proportion of females who are sexual or, equivalently, the level of investment in sexual reproduction if the same female is able to invest in both sexual and asexual reproduction.

In order to calculate the frequencies of different classes of offspring at *t*+1, we observe that for a given offspring, there are four options for its mother (Fig. 2, two first decision nodes). First, she can be sexual or asexual, with respective probabilities *p* and 1–*p*. Second, she can be mutant (*a*) or wild type (*A*) at locus L, with respective probabilities *x* and 1–*x*. *a* mothers have relative maternity propensities 1+*s*_f_ and 1+*s*_a_, for sexual and asexual females respectively, whereas *A* mothers have a relative propensity of 1, regardless whether the mother is sexual or not. Dividing by the mean fecundity 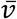 (the sum of all the numerators in the first two decision modes) gives probabilities that sum to 1. This becomes 1+*x*(*s*_a_+*ps*_f_– *xs*_a_), which reduces to 1+*xs*_f_ when *s*_f_=*s*_a_ (in scenario i), and 1+*pxs*_f_ when *s*_a_=0 (scenario iii). Intuitively, if *s*_f_ is negative (the *a* allele is deleterious), mean female fitness decreases with increasing *x* in both scenarios. In scenario iii (expression only in sexual females), mean female fitness also decreases with increasing *p*, and in a purely asexual generation (*p* = 0) mean fitness is 1. Note again that the above results are completely independent of *q* as long as 0<*q*<1 (with *q* = 1 the entire adult population is male, while with *q* = 0, females are only able to reproduce asexually).

**Figure 2:**
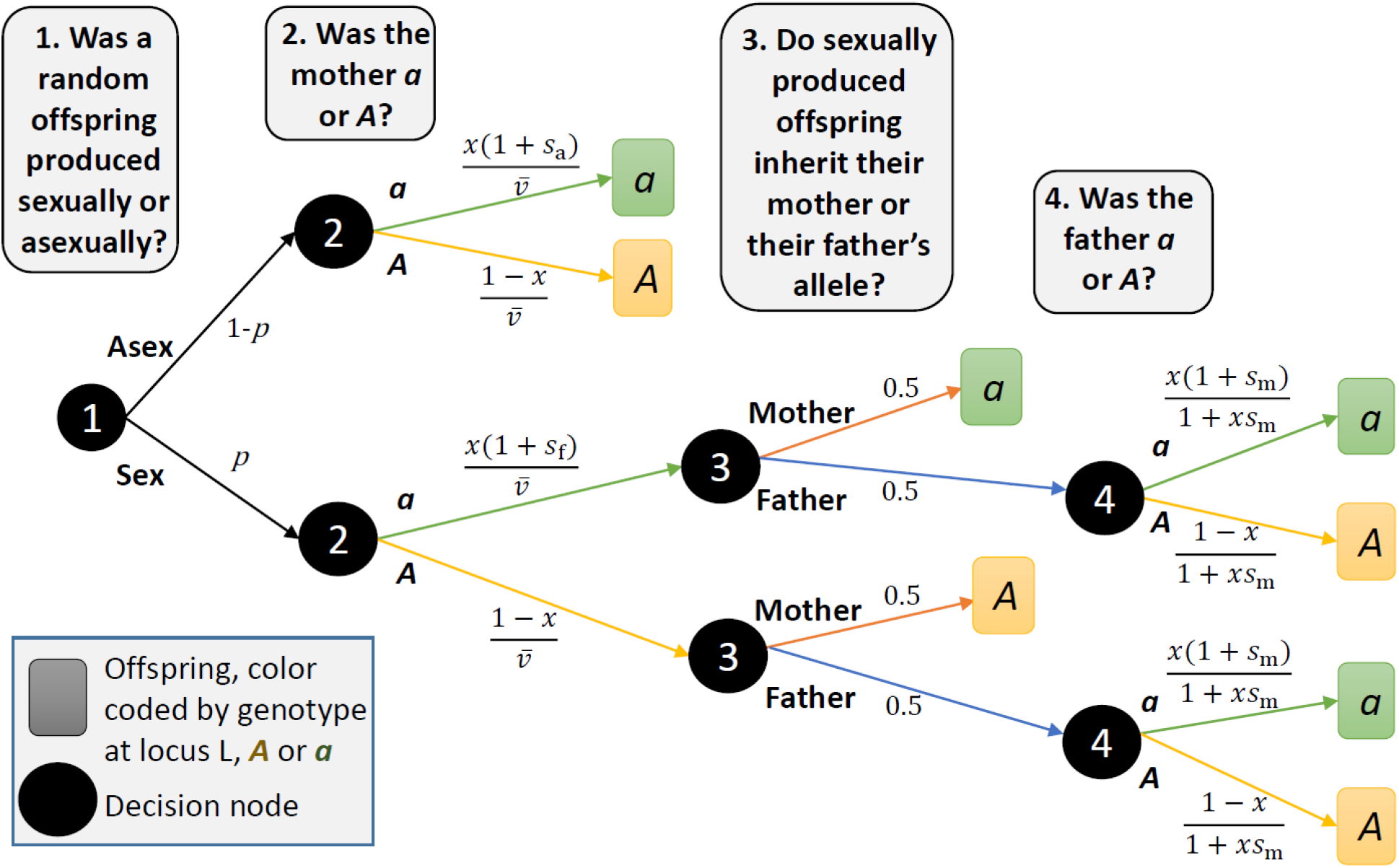
Decision tree to calculate frequencies of offspring produced by different classes of parents. Numbers along arrows indicate probabilities. Each node 1-4 (black circles) represents a dichotomous decision, and numbers along arrows indicate probabilities. Frequencies of each offspring class in the next generation are found by summing over all routes leading to a certain type of offspring (*A* or *a*, respectively yellow or green boxes), multiplying the probabilities along arrows leading to that offspring. Node 1: Each offspring can be produced sexually or asexually. Node 2: The mother can have genotype *a* or *A*. These two are the only relevant questions for asexually produced offspring, who copy their mother’s genotype. Node 3: Sexually produced offspring can inherit their mother’s or their father’s allele at L. This is the last relevant question for those inheriting their mother’s allele. Node 4: Offspring inheriting their father’s allele can have a father of genotype *a* or *A*.

The allele frequencies of asexually produced offspring are easy to derive, as they simply inherit their mother’s genotype. For sexually produced offspring, we assume Mendelian inheritance, i.e. half of the offspring inherit their mother’s genotype and the other half inherits their father’s genotype (Fig. 2, third decision node). For the sexually produced offspring inheriting their father’s genotype, the relative siring probabilities of the two types of fathers (*A* or *a*) are arrived at in the same way as for maternity (Fig. 2, fourth node). The genotype of the offspring with respect to locus L is shown in the colored squares at the end of each path in Figure 2.

Figure 2 allows us to find the total proportion of *a* offspring produced via all the different routes, by multiplying the probabilities along each route producing *a* offspring (green boxes), and summing over all the paths, Σ_*a*_. Starting from the top path (writing *x* for *x_t_* to simplify notation), we obtain:

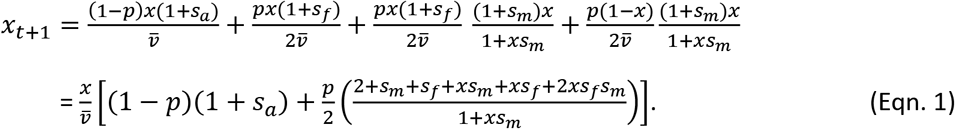

where 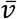 represents mean maternal fecundity, 1+*x*(*s*_a_+*ps*_f_–*ps*_a_).

### Calculating the realized selection coefficient in a sexual generation

Next we use Eqn. 1 to calculate the realized selection coefficient 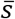 acting on a conditionally expressed allele during a sexual generation across the entire population, which permits a comparison with the results by Van Dyken and Wade (2010 p. 559). Specifically, we ask, what is the value of 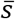 that, when applied to all individuals in a haploid population, yields the same change of allele frequencies as a given setting involving conditionally expressed alleles (with either males, all females, or sexual females expressing the relevant alleles)?

The change in mutant frequency in a generation where a proportion *p* of females reproduce sexually is given by Eqn. 1. We contrast this change with a constitutively expressed setting, where mutants have relative fitness 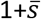 and 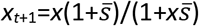. These two allele frequency changes are identical when

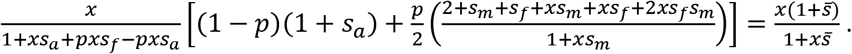

Solving for 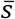 yields

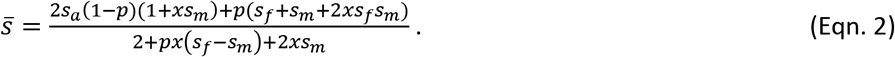

Checking some special cases to ensure that Eqn. 2 behaves sensibly, we see that when *s*_f_ = *s*_m_ = *s*_a_ = *s*, the expression simplifies to 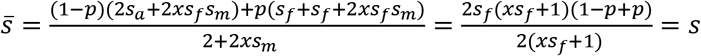, as it should.

Similarly, if we set *p* = 0, i.e. the population is fully asexual, we get

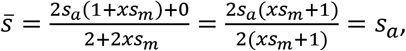

which is what we expect given that selection now only has its effect via asexual females.

For scenario (i), where only *s*_f_ ≠ 0, *p* = 0 yields 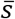 = 0, which is the correct value in this setting where all individuals are asexual and the locus is not expressed in asexuals. Additionally, setting *p* = 1 in Eqn. 2 makes *s*_a_ irrelevant, i.e. expression in asexuals does not matter when all reproduction is sexual.

Note that although selection coefficients *s*_m_ and *s*_f_ are not frequency-dependent, *x* generally does not cancel out in expressions for 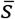 (Eqn. 2). The dependence of 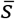 on *x_t_* is not very strong. It is negligible for small values of *s*, reflecting the assumption of weak selection in many population genetics analyses (e.g., Van Dyken and Wade (2010) assume that *s* is sufficiently small to justify ignoring terms involving *s*^2^), but the effect strengthens with *s*. In all our results, we use the full equations, and do not make any assumption that selection is weak.

## Results

### Change in genotype frequencies after one sexual generation

When locus *L* affects both asexual and sexual females (Fig. 3 and S1, top row), selection on males is overall less effective than selection on females: the effect of the male-specific selection coefficient *s*_m_ on the change in allele frequencies is weaker than that of the female-specific selection coefficients *s*_f_ = *s*_a_. Graphically, this is visible in Fig. 3 as allele frequency changes deviating much more from the *x*_*t*+1_ = *x_t_* diagonal when *s*_f_ is varied and *s*_m_ = 0 (top row) than when the opposite is true (middle row, Fig. 3; see Fig. S1 for an alternative graphical representation). While the deviations caused by selection on females become smaller as the proportion of offspring produced sexually (*p*) increases (right vs. left columns in Figs. 3 and S1), the effect caused by selection on males is opposite: the deviation grows with increasing *p*, reflecting males contributing more to the gene pool of the next generation when sexual reproduction is common. For expression in males only, or in sexual females only (middle and bottom rows of Fig. 3 and S1; scenarios ii and iii, respectively), the effect of selection on both sexes is very weak when *p* is low (deviations from the diagonal remain mild in Fig. 3C, E and S1C, E), but become stronger as *p* increases (Fig. 3D, F and S1D, F). As a whole (Fig. 3), the effects of selection are strongest (allele frequency changes are greatest) when selection acts on females across both reproductive modes (asex and sex), moderate when on males, and weakest when on sexual females only.

**Figure 3:**
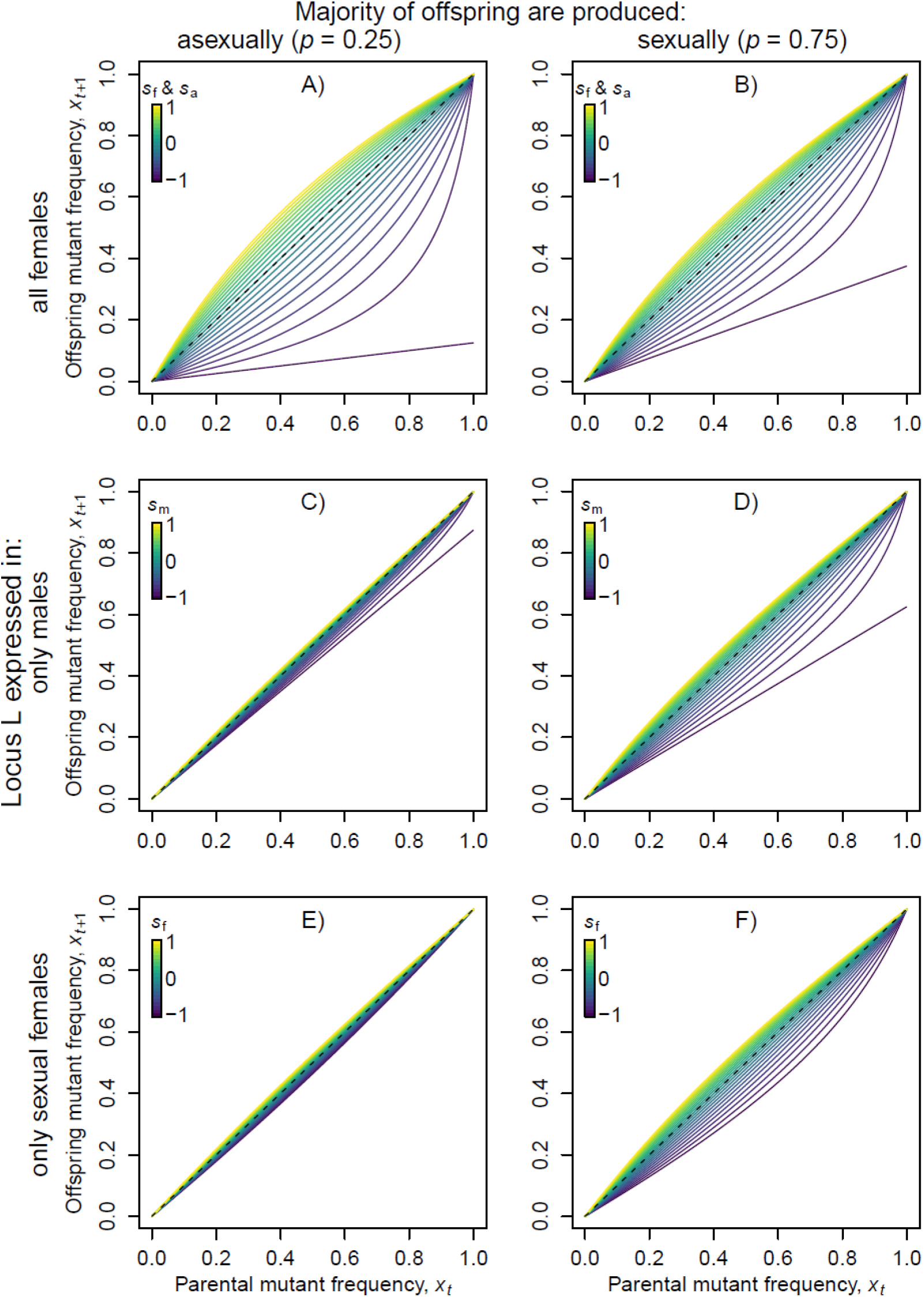
Frequency of mutant *a* individuals after one generation of reproduction, *x*_*t*+1_, for different parental mutant frequencies, *x*_t_. Thus, the stronger the deviation from the 1:1 line (dashed line), the stronger the effect of selection on allele frequencies. Columns: Majority of offspring produced (left) asexually vs. (right) sexually (*p*=0.25 vs 0.75). The selection coefficients in the expressing individuals are varied from 1 (yellow) to 0 (green) to −1 (purple), in steps of 0.1. Top row (scenario i): Expression in both sexual and asexual *a* females. *s*_m_=0, *s*_a_=*s*_f_ varied (line color). Middle row (scenario ii): Expression only in males. *s*_a_=*s*_f_=0, *s*_m_ varied (line color). Bottom row (scenario iii): Expression only in sexual females. *s*_a_=*s*_m_=0, *s*_f_ varied (line color).

Lethal alleles (*s*_m_, *s*_a_ or *s*_f_ equal to −1) show qualitatively different behaviour to non-lethal deleterious mutations across many scenarios. Eqn. 1 becomes linear on *x* for *s*_m_ = 0, *s*_f_ = *s*_a_ = −1 (scenario i) as well as for *s*_f_ = *s*_a_ = 0, *s*_m_ = −1 (scenario ii). However, it remains quadratic in scenario iii, i.e. for *s*_f_ = −1, *s*_m_ = 0 and *s*_a_ = 0 (see corresponding presence or absence of linear responses to *x* in the lowermost purple lines in Fig. 3). The disappearance of quadratic terms in scenario i reflects the fact that a female-expressed lethal *a* allele makes all females *A* in the relevant subpopulation. Therefore, *a* can only remain in the population via sexually reproducing *A* females whose offspring will be *a* with probability *x*/2, as the allele is not deleterious in males. If a proportion *p* of offspring are sexually produced, this means that the frequency of a lethal *a* drops from *x* to *px*/2 in a single generation, explaining the linear slopes in Fig. 3A-B. Similar reasoning applies for *s*_m_ = −1 (Fig. 3C-D), but now we assume that the lethal allele is not expressed in females, thus a proportion *x* is passed on unchanged via asexual females while it is halved in the proportion of the female population that reproduces sexually (since all sires are *A*), creating an overall milder drop in the frequency of lethal allele. Finally, in scenario iii, where lethality is only expressed if the female attempts a sexual cycle, *a* females are not available to mate with males of either genotype; if *a* is common, males have few females to mate with overall.

### Population-wide realized selection coefficients over a sexual generation

Above, we visualized allele frequency changes from one generation to the next. We next extend this to an analysis of 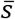 as a function of *s*_m_, *s*_f_, *s_a_* and *p* (Eqn. 2). Before proceeding, we set up some expectations, which are all based on the assumption that selection is ‘diluted’ whenever a parent escapes selection because of not expressing the locus in question. In an obligately sexual population where only one of the sexes expresses the locus (and the other has *s* = 0), then 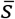 is half as strong as selection within the sex under selection, leading to a slope of ½ when graphing 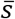 against *s*_f_ (or *s*_m_). Facultative sex can change this expectation by changing the patterns of dilution. If selection acts on female-expressed loci, and if all females express it (including asexuals, *s*_a_ = *s*_f_ ≠ 0, i.e. scenario i), then the slope is slightly shallower than the diagonal because some (but not all) offspring have fathers, and siring success is unrelated to male genotype. We expect the slope to be closer to ½ rather than 1, if sex is common. On the other hand, if only males (scenario ii) or only sexual females (scenario iii) express the gene, we expect the slope to be shallow overall. Dilution from asexual females, who do not express the *a* allele, is particularly strong if few offspring are produced sexually (low *p*).

These expectations are met by our model simulations (Fig. 4). In scenario i (expression in all females, top row of Fig. 4), selection is strongest (colored lines close to the 1:1 diagonal, i.e. 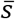 close to that of the female-specific selection coefficients) when a majority of offspring are produced asexually (Fig. 4A). Echoing the results of Fig. 3 A-B, we see that as the proportion of sexually produced offspring increases, the male contribution to genome of the offspring generation increases, leading to weaker selection (closer to the gray dashed line of slope ½, Fig. 4B). In scenario ii (expression in males only, middle row of Fig. 4) and iii (expression in sexual females only), selection is weaker overall (colored lines are flatter, i.e. the population-wide realized selection coefficient 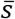 is closer to 0 for most *s*_m_), but strengthens as the proportion of sexually produced offspring increases (steeper slopes in Fig. 4D and F than in Fig. 4C and E).

**Figure 4:**
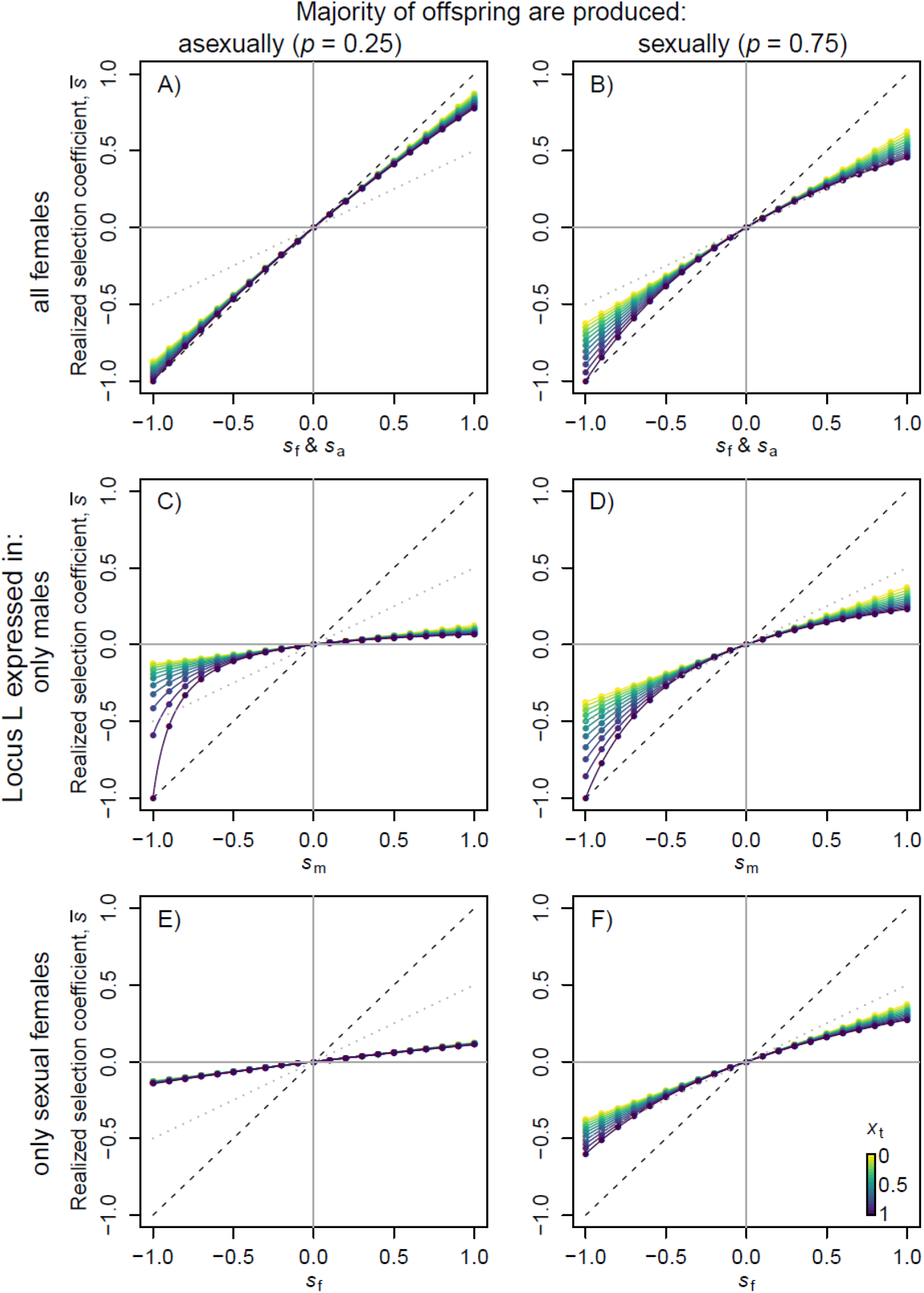
Population mean selection coefficient 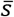 as a function of different sex-specific selection coefficients *s*_m_ and *s*_f_ for different values of *p* (columns) and mutant frequencies (*x_t_*, point color). Scenarios are the same as in corresponding panels in Figure 3. Dashed black 1:1 line indicates that the sex-specific selection coefficient equals the realized selection coefficient of the population. Dotted grey 1:2 line indicates the population realized selection coefficient that would be predicted from Van Dyken and Wade (2010) given an equal sex ratio (*q*=0.5), whereas our results hold for any 0<*q*<1 (but see Supplementary Material section 2). Note that in panel E, all lines and points are practically on top of each other, causing only the purple to be shown.

Additionally, the graphs make the emergent dependence on *x*_t_ clear: despite no *a priori* assumption that *s*_m_, *s*_f_ or *s*_a_ depend on *x*, the corresponding 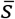 values indicate stronger negative selection when the deleterious alleles are at high frequency. Strongly deleterious selection coefficients (left end of each panel in Fig. 4) lead to stronger negative 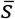 when *x_t_* is high (purple dots) than for medium and low *x_t_*. Conversely, beneficial mutations are most strongly selected for when they are rare (in upper right quadrants of each panel in Fig. 4, purple dots show weaker positive 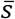 than green and yellow dots). The effect also strengthens with the absolute magnitude of *s* (i.e. in Fig. 4, the vertical spread of points/lines at a given location is always wider towards the edges of the plots).

### Comparison with *s*_c_ in Van Dyken & Wade (2010)

Next we compare the realized selection coefficients 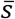 shown in Fig. 4 with those derived by Van Dyken and Wade (2010). Specifically, they define *s*_c_ as the “average selection against the conditionally expressed allele” (Van Dyken and Wade 2010, section “Expression in a fraction of individuals, *Φ*, at every generation”, p. 558). In a generation where a fraction *Φ* of the population expresses the conditionally expressed allele, they use the result *s*_c_ = *Φs* when the fitness of expressers is 1–*s* (note that a positive *s* in this formulation means that the mutation is deleterious; we take note of this notational difference when comparing our results to theirs).

In the specific context of facultative sex and sex-specific expression (“Expression in some individuals, some of the time”, p. 559), they interpret *Φ* as the sex ratio, expressed as the proportion of males in the population and denoted *Φ*_I_, where the I subscript refers to fraction of individuals expressing the trait (to emphasize the difference to *Φ*_G_ the fraction of generations expressing the trait). In our Fig. 4 their statement would show up as a linear relationship with *s*_m_, with a slope equal to the proportion of males *Φ*_I_. The difference between their prediction and ours remains moderate when the proportion of sexual females is high (right column of Fig. 4), but when asexuality is common (lower *p*), the difference becomes substantial.

We have shown that the fraction of expressers is not an adequate metric to identify the strength of selection when sex-specific selection is involved (Fig. 5). Specifically, the proportion of males *q* has no effect on 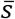 in our main model scenarios (but see Supplementary Material section 2); what matters for the contribution of males is *p*, the proportion of female reproductive investment allocated to sexual offspring (compare panels C and D, Fig. 4). Thus, for any proportion of males in the population, the population’s realized selection coefficient will vary widely depending on *p*: if a rare mutation (*x* near 0) is only expressed in males, the realized selection coefficient is approximately equal to *ps*_m_/2 (Eqn. 2 with *s*_f_ = *s*_a_ = 0 and terms including *x* considered small).

**Figure 5:**
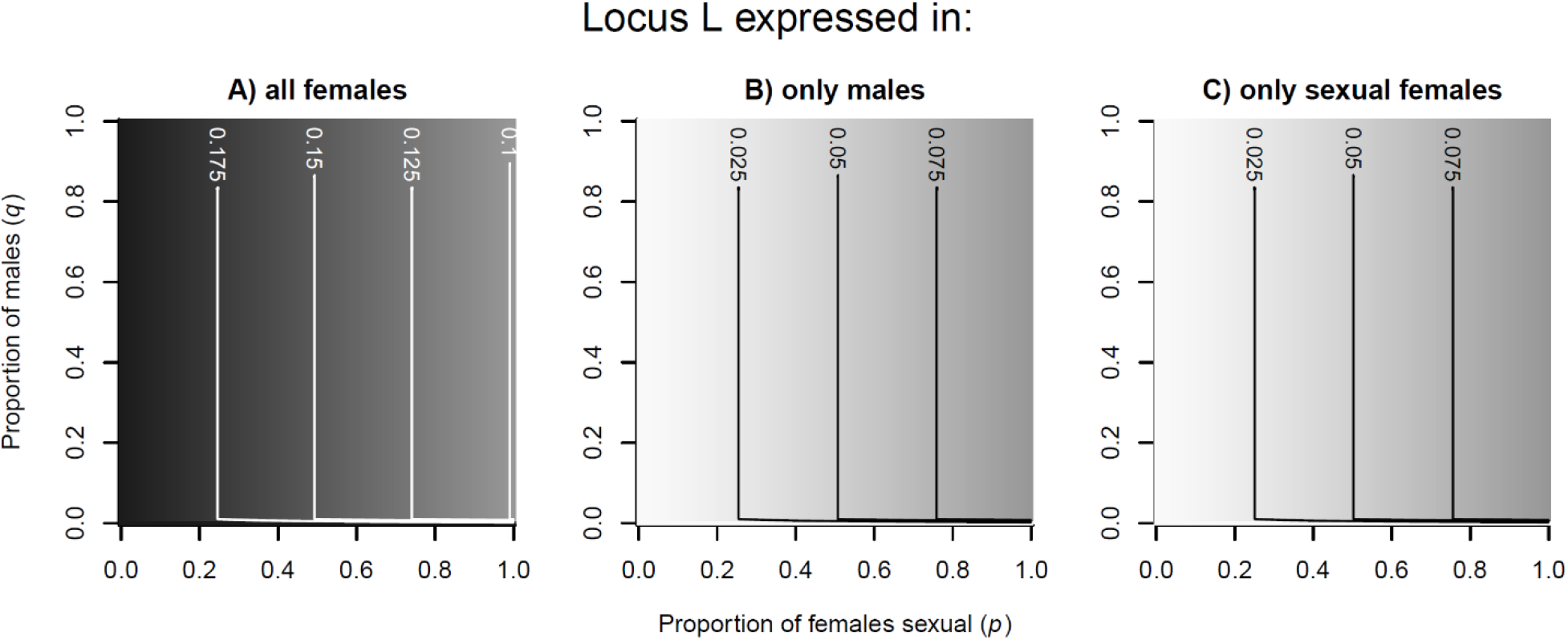
Heatmaps showing the realized selection coefficient (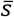) of the population as a function of the proportion of offspring produced sexually (*p*) and the proportion of males in the population (*q*). In all panels, *x*=0.1. A: Expression in both asexual and sexual females (scenario i), *s*_f_=*s*_a_=0.2 and *s*_m_=0. B: Expression only in males, *s*_f_=0 and *s*_m_=*s*_a_=0.2. C): Expression only in sexual females (scenario iii), *s*_f_=0.2 and *s*_m_=*s*_a_=0.

Our calculation of 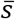 can be seen as a ‘plug-in’ to the model of Van Dyken & Wade (2010), in the sense that it can be used to replace their *s*_c_ = *Φ_s_* in all of their interesting analyses of the consequences of weakening selection strength on conditionally expressed loci. These include (but are not limited to) fixation probabilities for weakly deleterious or beneficial mutations, mutational load under mutation-selection balance, ratios of nonsynonymous to synonymous substitutions (dN/dS ratio), and rates of sequence divergence. Although we do not attempt to recreate all of their results here, we briefly focus on the effect of conditional expression on mutational load, adding our results to their Figure 1A as an illustrative case (Fig. 6). Van Dyken & Wade (2010) quantify this by ‘*R_p_*’, the ratio of the equilibrium frequency of a conditionally expressed gene to that of a constitutively expressed one (their equations 2 and 3). Rarity of expression associates with a higher load (higher frequency of deleterious mutations) in their model.

**Figure 6:**
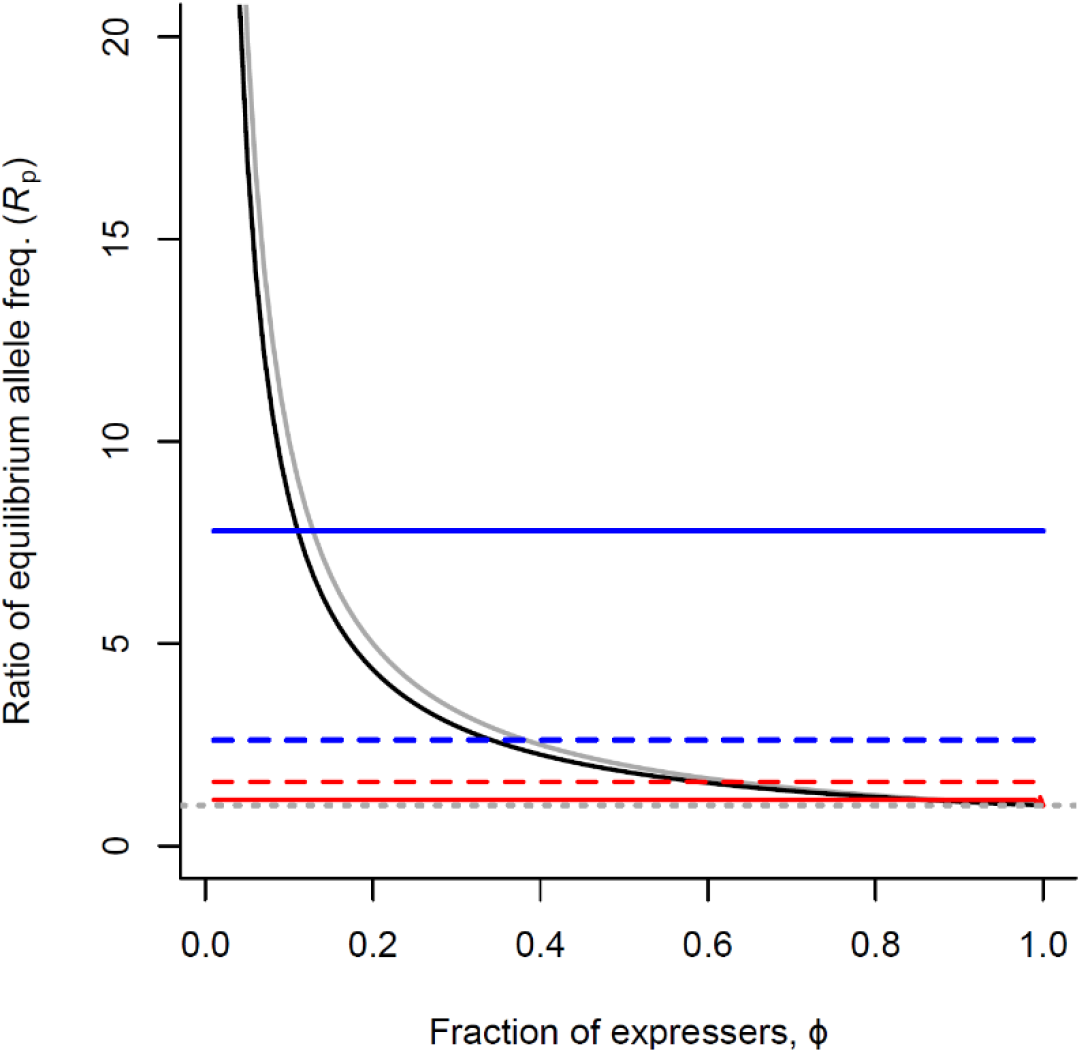
How conditional gene expression affects the equilibrium frequency of deleterious mutations (*s*=−0.3) at mutation-selection balance (modified from Van Dyken & Wade (2010); Fig. 1A). A ratio of *R*_p_ = 10 indicates that at a conditionally expressed locus the deleterious mutations are expected to have a 10 times higher equilibrium frequency than those at constitutively expressed loci. Grey solid line: An allele expressed by a proportion *Φ* of individuals in every generation (Van Dyken & Wade (2010); Eqn. 1). Black solid line: An allele expressed by all individuals in a proportion *Φ* of generations (Van Dyken & Wade (2010); Eqn. 2). Colored lines: Our result for alleles expressed only in females (scenario i, red) or only in males (scenario ii, blue) when *p*=0.25 (solid lines) or *p*=0.75 (dashed lines). The grey dotted line indicates *R*_p_=1, i.e. conditionally and constitutively expressed loci are equally likely to harbor deleterious mutations.

However, as a corollary of our prediction that sex ratio has no effect on 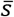, our model shows that it also has no effect on the level of deleterious mutations at mutation-selection balance (Fig. 6, but see Fig. S5 for non-baseline cases where selection coefficients themselves are a function of the sex ratio). What matters, once again, is relative female investment in sexual offspring: in our scenario i, male-expressed deleterious alleles reach 7.8 times the frequency of constitutively expressed alleles if female investment in sexual offspring is low (*p*=0.25, solid blue line), but this ratio drops to 2.6 as *p* increases to 0.75 (dashed blue line in Fig. 6). These ratios are much higher than those predicted by Van Dyken & Wade (2010) when males are common: e.g. at 50/50 sex ratio (*Φ* = 0.5), Van Dyken & Wade (2010)’s results (black and grey lines in Fig. 6) are far below our blue solid line and somewhat below our blue dashed line). In contrast, when males are rare (e.g. *Φ* < 0.1) Van Dyken & Wade (2010) predict much higher levels of male-expressed deleterious alleles than we do, because they did not consider the Fisher condition which ensures that each male gains higher paternity when males are rare.

For female-specific loci, on the other hand, selection is almost as efficient as for constitutively expressed loci at low *p* (only a 14 % increase in deleterious mutations if *p*=0.25; solid red line in Fig. 6), but becomes less efficient as *p* increases and the male contribution to offspring genotypes increases (58 % increase in deleterious mutations if *p*=0.75; dashed red line in Fig. 6). If all reproduction is sexual, loci with sex-limited expression are expected to harbor twice as many deleterious mutations as constitutively expressed loci, be it in males or females, and regardless of population sex ratio, since all offspring have both a mother and a father (Fisher 1930).

In conclusion, our main finding that the proportion of individuals in the population (*q* or 1–*q* for males or females, respectively) ceases to be a good predictor of the efficiency of selection on a trait, holds true in our baseline model (scenarios i-iii). However, it is possible to construct scenarios where *q* matters alongside *p*; we explore some of these more special cases in the Supplementary Material section 2.

## Discussion

A substantial proportion of genes is only expressed in a fraction of population members or in some generations. One of the most obvious cases of such conditional gene expression are the numerous genes with male and female specific expression (Mank 2017; Molinier et al. 2018). In many situations (but not all, see Harts et al. 2014), sex-specificity translates into a simple expectation of a ‘tug-of-war’ where the overall selection coefficient can be computed by averaging over the sex-specific effects. Here we show deviations from this simple expectation when sex is facultative. Specifically, the realized strength of selection 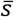 acting on genes that are only expressed in a fraction of the population cannot be predicted simply by the fraction of expressers, *q*, when these expressers are males. In other words, the general theory of conditional gene expression proposed by Van Dyken & Wade (2010) does not straightforwardly apply to cases where only some offspring have a father.

Our modelling explicitly takes into account the Fisher condition (every fertilized egg must have a father; Queller 1997; Houston and McNamara 2005) and found it to be responsible for breaking the link between the frequency of expressers and selection, when the expressers are males of facultatively sexual species. For females, however, the gist of Van Dyken & Wade (2010) remains valid: selection remains stronger overall if female genotype impacts fitness regardless of the current reproductive mode of the female, than if females only express the relevant alleles when reproducing sexually.

We note that Van Dyken & Wade (2010)’s predictions for mutation accumulation and sequence divergence for genes with sex-limited expression still hold true in obligate sexuals: selection strength is halved relative to genes that are expressed in both sexes (Charlesworth et al. 1987; Whitlock and Wade 1995; Barker et al. 2005). This also appears as a special case in our models: when *p*=1, 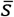 reduces to *s*/2 for mutations at low frequency (if we let terms including *x* become 0) in all model scenarios (Eqn. 2, S2, S5). However, since Van Dyken & Wade (2010)’s suggestion that rarity of males *per se* can account for weakened selection on male-specific genes does not hold, interpretations of empirical data need to be re-evaluated. A good example is their evaluation of Brisson & Nuzhdin (2008)’s study of pea aphids *Acyrthosiphon pisum*, where male-specific genes have approximately twice as high levels of sequence divergence in among-population comparisons as female-specific or sex-neutral genes. Although this pattern is qualitatively in agreement with frequency-based theory, the difference is an order of magnitude weaker than Van Dyken & Wade (2010) predict based on male rarity. Male aphids appear only at the end of summer, after 10 to 20 generations of parthenogenesis, giving *Φ*_I_ between 0.05 and 0.1. Since all offspring that survive the winter have necessarily been produced sexually (*p*=1, or similarly *C*_a_=1, see Supplementary Material section 2a), *R*_p_ in this system would amount to between 20 and 40. Thus, the pea aphid data are in much closer agreement with our predictions of a more moderate effect of sex-specific gene expression.

Biological interpretation of the parameter *p* in our model is not necessarily limited to the proportion of offspring produced sexually. An equally interesting application is to species where hermaphrodites coexist with unisexual males, as in *Caenorhabditis elegans* (Hodgkin and Doniach 1997; Chasnov and Chow 2002). Van Dyken & Wade (2010) explain the poorer mating success of *C. elegans* males relative to that of males in the obligately sexual sister species *C. remaneii* through the rarity of unisexual males (approximately 5 % of individuals every 5 generations) causing accumulation of deleterious mutations (*R*_p_ = 1/*Φ*_I_*Φ*_s_ = 1/(0.05×0.2) = 1/100). If we define *p* as the proportion of offspring fathered by unisexual males, our model shows that this result is valid only if females mate with unisexual males (rather than self-fertilizing or mating with another hermaphrodite) in approximately the frequency predicted by male density under random mating. Random mating across all sex ratios seems *a priori* unlikely given that many hermaphrodite species avoid selfing, but also have better access to themselves than to other individuals (including males) should mate encounters become difficult. Therefore, traits that affect male competitive success when females (or the female function of hermaphrodites) mate multiply, such as the production of copulatory plugs in *C. elegans* (Hodgkin and Doniach 1997), may be key to securing matings for unisexual males (see Supplementary Material section 2b). This can prevent mutational degradation of the male genome (Chasnov and Chow 2002). Testing the predictions of our model quantitatively will require measuring the proportions of offspring fathered by unisexual males versus hermaphrodites, which is rarely done in natural populations of *C. elegans* (Félix and Braendle 2010).

While our main results show why there is a ‘null expectation’ that the strength of selection is independent of the sex ratio, we also consider cases where additional processes come into play and make selection dependent on the sex ratio (Supplementary material section 2). One category of such a causality (Supplementary material section 2a) assumes that the reproductive success of females using either sex or asex as their current reproductive mode depends on the sex ratio, e.g. because one of the two reproductive modes experiences more male harassment, and harassment is more intense when males are common (Long et al. 2009; Lankinen and Karlsson Green 2015; Kobayashi 2019). Low fecundity of sexual females at very low mate availability is another possibility within this category, either because of actual matelessness (e.g. Kiørboe 2007; Rhainds 2019) or because of time, energy or predation costs of mate-searching. Another category assumes that the selection coefficients *s*_a_, *s*_f_ and *s*_m_ themselves become a function of *q*, and in our Supplementary material section 2b we analyze a case where the strength of selection on males depends on *q* because females mate predominantly monandrously when males are scarce, and polyandrously when they are common (Duneau et al. 2020); therefore, selection on males to succeed in sperm competition becomes stronger with male frequency. These additional analyses add nuance to our baseline model: it is possible to find causal routes where *q* can alter 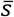, but this requires making additional and system-specific assumptions, and so these do not act to restore any simple expectation that 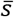 is proportional to the frequency of expressers.

### Conclusion

Modern lab techniques, such as RNAseq, have changed our understanding of gene expression in different life stages and conditions, and have revealed a large numbers of genes with differential expression in males and females. It is now possible to test whether these genes differ in their genomic signature from genes assumed to have sex-independent expression. Such differences are expected to come about because selection on these genes is on average weaker, and the effects of genetic drift thus become more apparent, as compared to genes expressed in all members of a population. These genes are therefore prone to deleterious mutations, that are less likely to become purged from the population, and beneficial mutations may be more easily lost before they can go to fixation. This will result in conditionally expressed genes having, among other population genomic summary statistics, higher ratios of non-synonymous to synonymous mutations (dN/dS). Predictions made by mathematical modelling of these processes can now be tested with real data, and vice versa, findings from genomic analysis can be placed in a proper theoretical framework.

Our study provides a theoretical backbone for predictions regarding the efficiency of selection on genes with sex-limited expression in facultatively sexual organisms such as many plants, Cladocera (water fleas), many insects (e.g. aphids), rotifers, cnidarians, and bryozoans. The former predictions for sex-specific genes in such organisms predicted too strong effects of rare sex, i.e. overestimating the reduction in the efficiency of selection, especially in males. Our model shows that these effects are more modest and that therefore the genomic signatures of sex-specific genes differ less among sexes and differ less among genes with sex-specific expression and constitutively expressed genes than previously predicted. Without the effect of the Fisher condition constraining mutational meltdown on genes with sex-limited expression when sex is rare, as in cyclical parthenogens and many other facultatively sexual organisms, this widespread reproductive mode might even have been impossible to maintain.

## Acknowledgments

We thank David Duneau for discussions motivating this work. DE and HK are supported by the Swiss National Science Foundation.

## Author contributions

All authors initiated the project. TRH and HK developed and analyzed the models, with input from DE. TRH wrote the manuscript with input from all authors.

## Supplementary Material

### Supplementary Material contents

#### 1. Plots of Δ*x*

**Figure S1:**
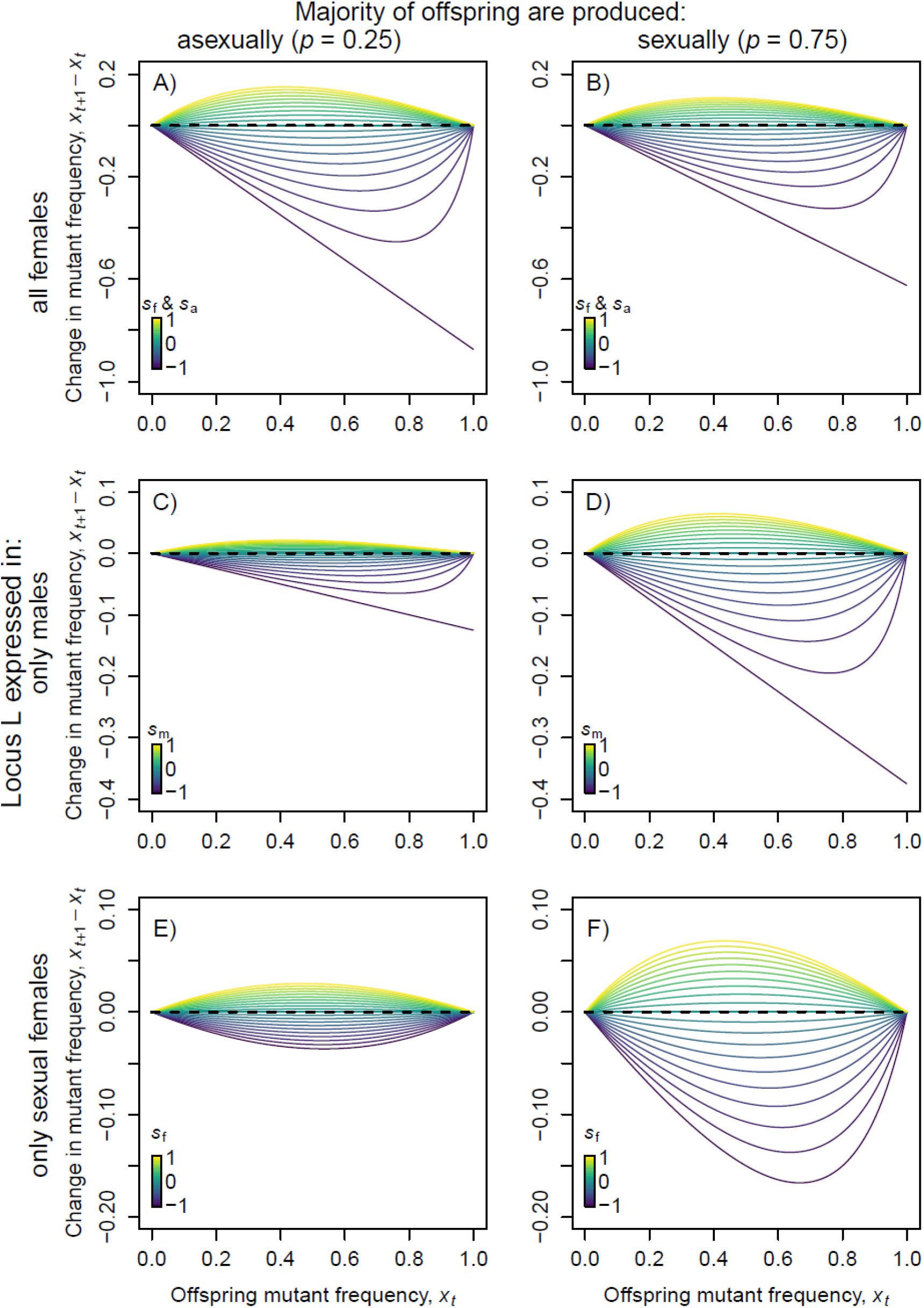
Equivalent to Fig. 3 in the main text, except plotting Δ*x* = *x*_*t*+1_-*x_t_* over *x_t_* rather than *x*_*t*+1_ over *x_t_*. Note the difference in y-axes across rows.

#### 2. Model extensions

Here we expand our baseline model by allowing sexual and asexual female to differ in their reproductive success in a manner that depends on the sex ratio (section a) and by allowing selection on traits expressed by a specific category of individuals to depend on q, which we exemplify by male-male competition that operates differently depending on sex ratio (section b).

##### a. Sex ratio modulates the reproductive success of sexual and asexual females

Costs of the two modes of reproduction (sexual and asexual) may plausibly depend on population sex ratio, *q*, since male presence may have negative impacts on female fitness but, for sexual females, very low male availability may also be detrimental. We model such differences as costs associated with either reproductive mode, which can be interpreted as modifications of either maternal fecundity or juvenile survival (their effects are identical in our model that has discrete generations and density-independent fecundity). We use *c*_s_ and *c*_a_ (subscripts ‘s’ and ‘a’ refer to respectively sexual and asexual females) to quantify the reduction in the propensity of sexual or asexual females to be the mother of a randomly chosen offspring. The cost terms range between 0 (no cost) to 1 (the mode of reproduction is impossible). The maternity propensities for each class of mother (asexual or sexual, wild type *A* or mutant *a* at locus L) become

- Asexual, *a*: (1–*p*)*x*(1+*s*_a_)(1–*c*_a_)
- Asexual, *A*: (1–*p*)(1–*x*)(1–*c*_a_)
- Sexual, *a*: *px*(1+*s*_f_)(1–*c*_s_)
- Sexual, *A*: *p*(1–*x*)(1–*c*_s_).

Summing these values to obtain normalized probabilities of maternity we arrive at a mean maternal fitness (equivalent to the denominators at decision 2, Fig. 2),

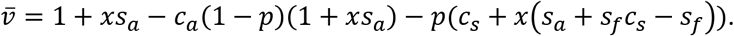

This expression leads to some intuitive results. If both cost terms are set to 0, mean maternal fitness shortens to those used in Fig. 2 (decision 2). As *c*_s_ increases towards 1, mean fitness decreases, at a rate scaling with *p*, the frequency of sexual reproduction. If *p* = 1 (all females are obligate sexual) and *c*_s_=1 (sexual reproduction is impossible), the population is obviously unviable and 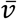 shortens to 0. The same occurs if *p*=0 and *c*_a_=1, as it should: A population reproducing entirely asexually goes extinct if asexually produced offspring are unable to survive.

We now turn our attention to the allele frequencies at *L* in the next generation, *x*_*t*+1_, and the realized strength of selection, 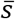. Using the above probabilities of maternity as in the main text (probabilities of paternity remain unchanged since all males are sexual), we obtain

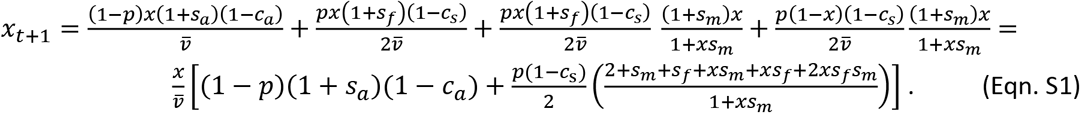

Again as in the main text, we find the population realized strength of selection 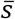 by comparing Eqn. S1 to the allelic frequency change that occurs in a constitutively expressed locus where mutants have relative fitness 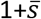, which is 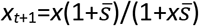. These two allele frequency changes are equal when

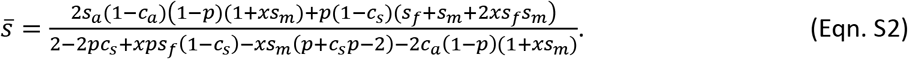

Although these equations for *x*_*t*+1_ and 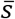 look somewhat unwieldy, they behave as they should and shorten to the expressions for the main model (Eqns. 1 and 2) when *c*_s_ and *c*_a_ are both zero. We also note that all terms containing a cost of sex become 0 when *p*=0, as expected – i.e. when all reproduction is sexual, costs of sex do not matter for selection.

We now look at some simple examples of how the density of males, *q*, can affect selection on conditionally expressed loci by altering the relative reproductive success of sexual versus asexual females.

First, consider a case where male harassment affects sexual females negatively, while asexuals avoid being harassed entirely. Costs to sexual females plausibly depend on both *p* (proportion of females that are sexual) and *q*. We examine the function

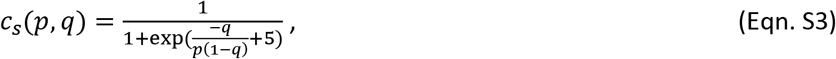

which is an increasing function of the number of males, *q*, per sexual female, *p*(1–*q*). The function is of a logistic shape (Fig. S2) to bound it between 0 and 1.

Since the cost *c*_s_ now depends on *q*, selection as a whole does so as well (Fig. S3). A high ratio of males to sexual females harms offspring production via the sexual route. Fewer offspring have a sexual mother when harassment impacts such mothers negatively. The outcome of harassment costs therefore recapitulates the general patterns associated with low *p* in the main text: The strength of selection decreases for loci expressed only in males (scenario ii, Fig. 5B) or only in sexual females (scenario iii, Fig. 5C), but increases for loci expressed in both asexual and sexual females (scenario i, Fig. 5A). Here, in the extreme case where sexual reproduction becomes impossible, 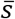 becomes identical with the selection coefficient acting on asexual females (*s*_a_=0.2) in scenario i (Fig. S3A), and takes the value 0 in scenarios ii (Fig. S3B) and iii (Fig. S3C). These patterns are familiar from the model in the main text, where, however, the effect of *p* was not modulated by any variation in the value of *q*. In the present version, changes in *q* can affect the realized strength of selection on conditionally expressed loci by altering the success of sexuals relative to asexuals in contributing to the next generation.

**Figure S2:**
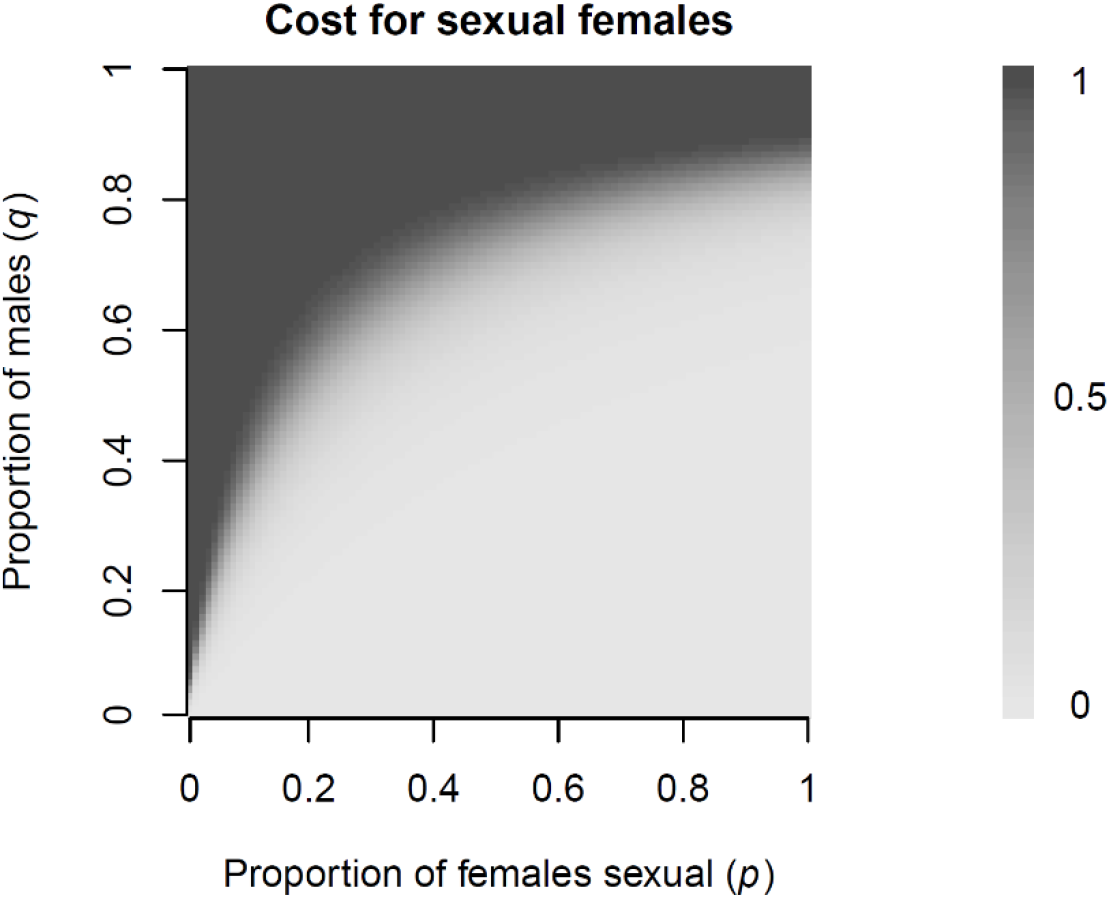
Heatmap showing cost to sexual females *c*_s_ as a function of population sex ratio (proportion of males, *q*) and the proportion of females that are sexual, *p*.

**Figure S3:**
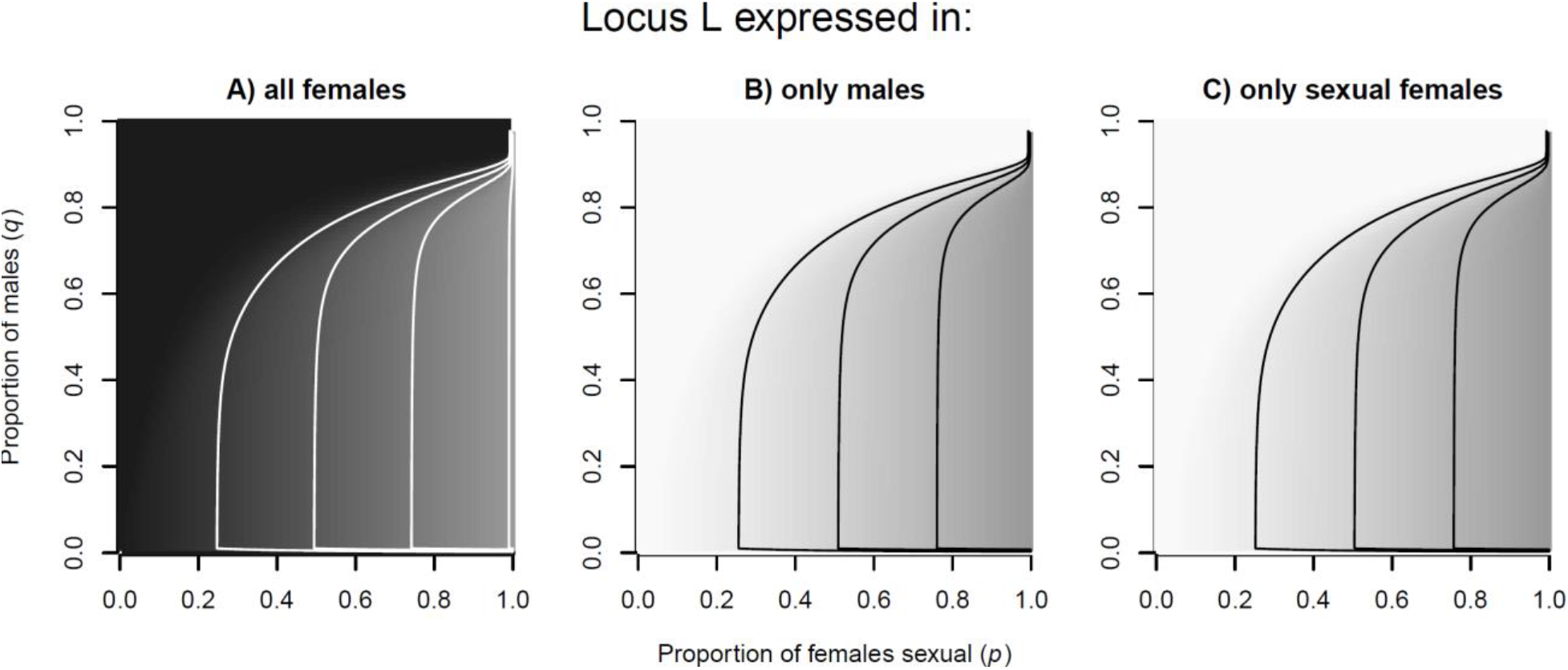
Heatmaps showing the population mean selection coefficient (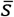) as a function of the proportion of offspring produced sexually (*p*) and the proportion of males in the population (*q*), when males exert costs on sexual females. In all panels, *x*=0.1. A: *s*_f_=*s*_a_=0.2 and *s*_m_=0 (scenario i). B: *s*_f_=*s*_a_=0 and *s*_m_=0.2 (scenario ii). C: *s*_m_=*s*_a_=0 and *s*_f_=0.2 (scenario iii). Shades span from 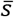=0.2 (black) to 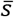=0 (white). From left to right, contour lines show 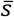 values of (in A) 0.175, 0.15 and 0.125; and (in B and C) 0.025, 0.05 and 0.075.

It is also possible to envision a range of other effects of male density *q* on the fitness costs of sexual or asexual females. Since the results are intuitive, we list them briefly without producing figures for them. Male harassment may not be limited to sexual females; it may even be more damaging to asexual females (who resist matings, Gerber and Kokko 2016) than sexual females (who accept them). This can either lead to damage to the asexual female, or (at least in organisms where a given female can reproduce both sexually and asexually) to the asexual female ‘giving in’ and reproducing sexually instead (Dagg 2006). Both possibilities decrease the probability that a given offspring in the next generation was produced asexually, leading to stronger selection on loci expressed only in males or sexual females, and weaker selection on loci expressed in all females (since the lower success of asexual females leads to a larger genetic contribution from males). The intensity of such costs to asexual females can be expressed using a similar function as that for *c*_s_(*p*, *q*) above (Eqn. S3), except that costs now increase as a function of 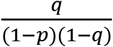, i.e. costs become more intense as asexual females become fewer.

Of course, it is also possible that harassing males are unable to discriminate between sexually and asexually reproducing females, or their presence imposes costs that are felt in local depletion of resources (food consumed by males is equally unavailable to sexual and asexual females). Costs that are felt equally by sexual and asexual females can be modelled as both *c*_s_ and *c*_a_ becoming a function of *q*(1-*q*). In this case, the relative importance of sexual and asexual reproduction in producing the next generation are unchanged, and the effect of *q* disappears.

Finally, scarcity of males may also impose costs on female reproduction, if sexual females become sperm-limited and unable to reproduce. Infertility risk and/or mate-searching costs impact sexual but not asexual females, and this means that *low* q associates with similar outcomes as *high q* in Fig. S3 (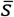 increases in scenario i, and decreases in scenario ii and iii).

##### b. Selection on traits depends on the sex ratio: an example with male-male competition

In the baseline model, we assumed that category-specific selection coefficients *s*_a_, *s*_f_ and *s*_m_ did not depend on either *p* or *q* (while we did proceed to investigate the demographic consequences of p and q that then determine 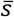. In real life, this independence assumption may be violated. We exemplify such a violation in the context of *s*_m_ depending on *q*. Traits with male-limited expression may impact a male’s attractiveness or competitive ability. The relationship between trait value and male mating success may depend on *q*. To provide an illustrative example, we present a simple model where the genotype at locus *L* determines a male’s success in sperm competition.

Assume that females mate with either 1 or 2 males (this range captures the overwhelming majority of matings in sexual generations of the cyclical parthenogens *Daphnia magna*, Duneau et al. 2020). We assume that polyandrous matings occur with a probability *f*(*q*), a function that increases with *q*. In monandrous matings, male genotype does not matter, since the male fertilizes all eggs of the female with probability 1, whereas in polyandrous matings, *A* and *a* males have relative competitive successes 1 and 1+*s*_0_, respectively. We denote the difference in competitiveness by *s*_0_ rather than *s*_m_ here, since it may not reflect the overall selection strength on *L* (in monandrous matings, genotype does not affect fertilization success).

If *x* is the proportion of males with genotype *a*, the chances of an egg being fertilized by an *a* male is

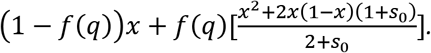

In other words, in a polyandrous mating, the probability that both males are *a* is *x*^2^, and the probability that only one of the males is *a* is 2*x*(1-*x*). Only in this second case does difference in sperm competitive ability play a role. (With probability (1-*x*)^2^ neither male is *a*, a term that disappears since it is multiplied by 0.)

Thus, the actual selection coefficient, *s*_m_, may be different from that predicted by the relative competitive abilities *s*_0_, depending on *q*. To find what *s*_m_ results from a given *q*, we observe (from Figure 2, fourth decision node) that as a whole, an egg is fertilized by an *a* male with probability *x*(1+*s*_m_)/(1+*xs*_m_). We are then interested in solving

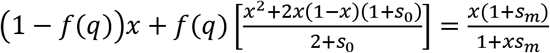

for *s*_m_, which yields

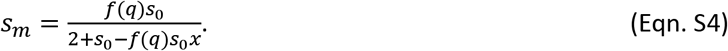

This shows how effective selection strength depends on *q* (via *f*(*q*)), and also positively on *x*. As an example, assume that the probability of polyandrous matings is a power function of *q*, denoted *q^k^*, which can increase in the [0,1] range either steeply from 0 (if *k*<1) or slowly, only rising to near 1 if *q* becomes very high (for *k*>1). Then effective selection strength becomes 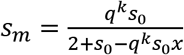, which always leads to *s*_m_<*s*_0_, i.e. effective selection on males *s*_m_ is weaker than the differences in competitive ability (Fig. S4).

As the frequency *q* of males in the population increases (and thus also the probability of polyandry, *f*(*q*)=*q*^k^), effective selection on males increases, but is still considerably weaker than *s*_0_, only approaching *s*_0_/2 for high *x*. Thus, this effect of sex ratio can produce substantial *q*-dependence of selection strength at the population level 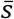, leading to 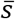 near 0, especially for high k (Fig. S5B). When *k*=2, the effect of sex ratio *q* is of comparable strength to that of *p*, the proportional female investment in sexual offspring (Fig. S5B).

The potential for monandry to weaken selection on males becomes clear when looking at accumulation of deleterious mutations in males (Fig. S6), as we do in the main text in Fig. 6 (comparing our results to Van Dyken & Wade (2010); Fig. 1A). For low *k* (blue lines), which means that polyandry is frequent except when males are very scarce, the ratio of deleterious mutations at mutation-selection balance in male-limited versus constitutively expressed loci (*R*_p_) are lower than those found in Van Dyken & Wade (2010) at low male densities, and about one order of magnitude higher at high male densities. Increasing *k* (green lines) reduced polyandry across all sex ratios, and the outcome is logical: selection on males is much weaker and deleterious mutation accumulate at drastically higher rates. When males are rare enough for polyandry to be very uncommon, *R*_p_ skyrockets: when *p*=0.25 and *k*=2 (continuation of the solid green line in Fig. S6), we find R_p_=1360 and 1360,000 for *q*=0.1 and *q*=0.01, respectively.

Obviously, if polyandry is fairly frequent no matter what the value of *q*, the effect of *q* becomes milder (less change in *s*_m_ and 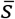 with *q*, with selection closer to *s*_0_/2). The same applies if polyandry was rare no matter what the value of *q*: overall weak selection with little dependence on *q*.

**Figure S4:**
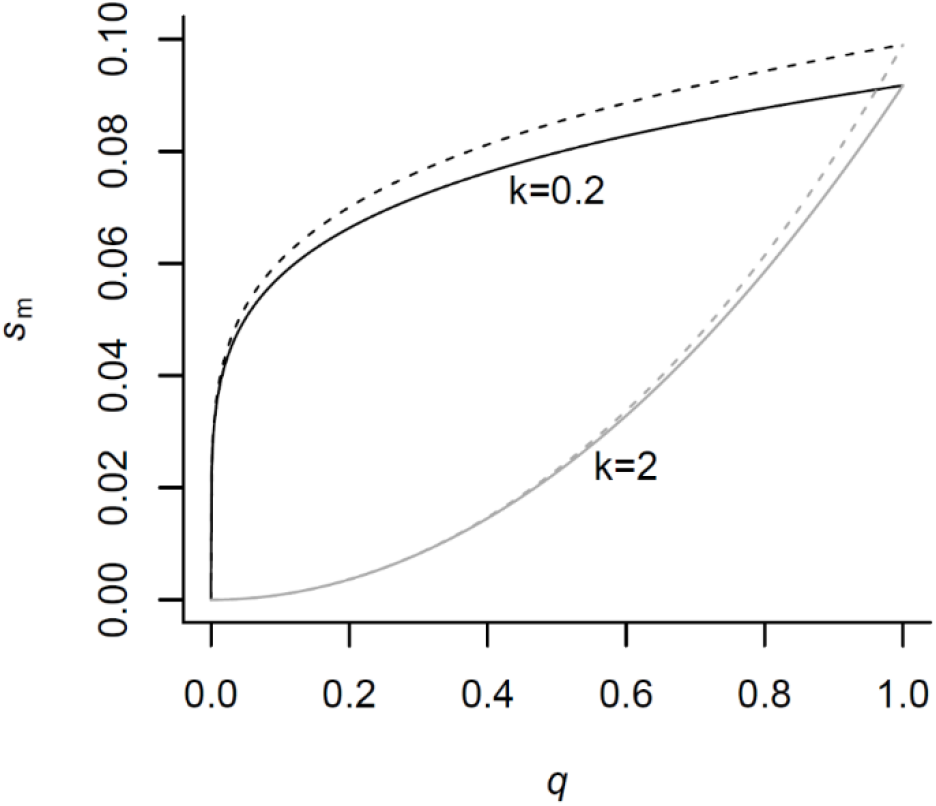
Effect of population sex ratio (*q*, proportion of males) on effective selection strength *s*_m_, when the frequency of polyandrous matings is *f*(*q*)=*q*^k^. Here, *s*_0_=0.2, and results are shown for *x*=0.1 (solid lines) and *x*=0.9 (dashed lines), and k=0.2 (black lines) and k=2 (grey lines).

**Figure S5:**
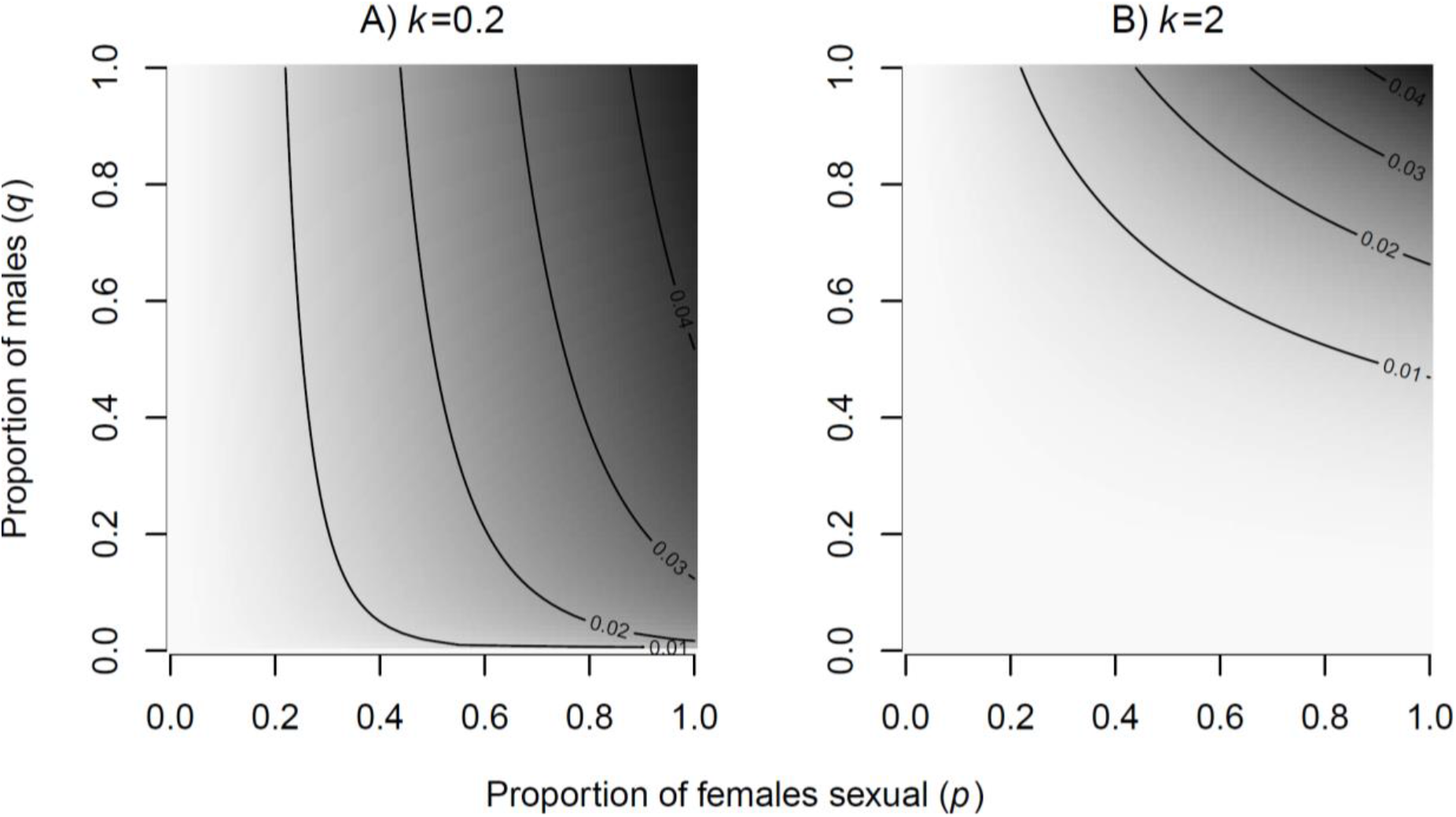
Heatmaps showing the population realized selection coefficient (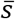) as a function of the proportion of offspring produced sexually (*p*) and the proportion of males in the population (*q*), when the locus *L* determines male mating success in polyandrous matings (e.g. sperm competition). In both panels, *x*=0.9, *s*_0_=0.2 and *s*_a_=*s*_f_=0. Results shown for when the frequency of polyandrous matings increases steeply (panel A, *k*=0.2) or slowly (panel B, *k*=2) with *q*.

**Figure S6:**
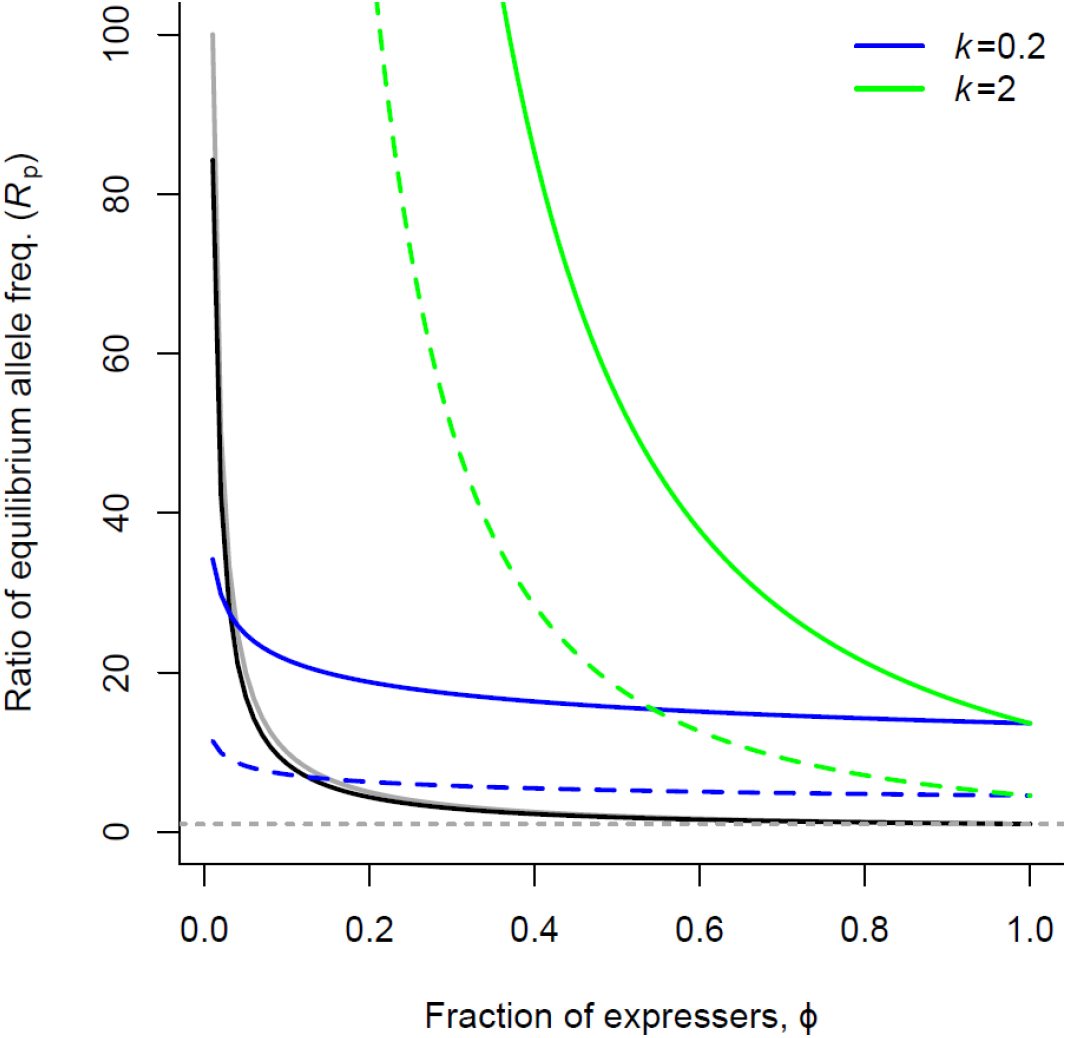
Accumulation of deleterious mutations (*s*=−0.3) on male-limited versus constitutively expressed loci at mutation-selection balance. E.g., a ratio R_p_=10 indicates that male-limited loci are expected to harbour 10 times more deleterious mutations than constitutively expressed loci. Colored lines show our results (‘Sperm Competition’ model) when the probability of polyandrous matings depends on male density (*q*, or *Φ*) according to a power function with parameter *k* equal to 0.2 (blue) or 2 (green), when a proportion *p*=0.25 (solid lines) or *p*=0.75 (dashed lines) of offspring are sexually produced. Gray and black solid lines are for comparison with Van Dyken & Wade 2010; Fig. 1 A), as in our Fig. 6. The gray dotted line indicates *R*_p_=1, i.e. conditionally and constitutively expressed loci are equally likely to harbor deleterious mutations.

